# Prediction of early *Toxoplasma gondii* infection during pregnancy combining host biomarkers and machine learning

**DOI:** 10.64898/2026.10.28.762409

**Authors:** Natalia Velasco-Jiménez, Roberto Sánchez-Sánchez, Ana Huertas-López, Camila Peres-Rubio, Francisco Huertas-López, Julio Benavides, Daniel Gutiérrez-Expósito, José Joaquín Cerón, Pilar Horcajo, Rafael Calero-Bernal, Luis Miguel Ortega-Mora

## Abstract

*Toxoplasma gondii* is a major cause of reproductive failure and severe gestational complications in both humans and small ruminants and early diagnosis is essential to prevent adverse outcomes. In sheep, congenital toxoplasmosis can trigger early fetal death (EFD) during the acute phase of the infection (days 7-14 post-infection) when conventional diagnosis methods are ineffective. Therefore, this study aimed to identify early infection biomarkers using a machine learning approach to support EFD diagnosis and providing a translational foundation for early diagnostic strategies in human toxoplasmosis. For that, 27 serum biomarkers (cytokines, acute-phase proteins, oxidative stress and hemostatic parameters) were evaluated during the early stage of infection in 22 pregnant sheep experimentally challenged with *T. gondii* and 6 uninfected controls. To assess diagnostic specificity, models were tested using sera from animals experimentally infected with other abortifacient agents (*Neospora caninum* and *Chlamydia abortus*) and validated with samples from 19 natural abortion outbreaks. A four-biomarkers panel, encompassing CXCL9, CXCL10, IFN-γ, and serum amyloid A, achieved over 74% accuracy in the specific detection of early *T. gondii* infection and 100% of sensitivity when studying natural infection cases. This approach might support the development of rapid diagnostic strategies, improving early intervention and disease management in livestock production systems. Its translation to human medicine could also aid in addressing current diagnostic gaps and facilitate timely interventions to prevent congenital toxoplasmosis and pregnancy loss in women.

**Importance:** This study addresses a critical veterinary and public health challenge: the diagnostic gap in early infection during congenital toxoplasmosis. In sheep, early fetal death (EFD) is a clinical outcome than remain underdiagnosed under field conditions due to the absence of maternal IgG antibodies and placental or fetal parasite DNA at the time of abortion. By pioneering the integration of host-response biomarkers with machine learning, this work successfully predicted acute infection with 100% diagnostic sensitivity and 85% accuracy under real-world field conditions (natural infections). Furthermore, this study provides clues for unraveling the pathogenic mechanism involved by demonstrating that EFD is driven mainly by oxidative processes rather than by the initially proposed increase in pro-inflammatory cytokines. In addition, the proposed approach may have a translational potential, both in terms of the selected biomarkers and the machine learning-based methodology, with the purpose of enabling early therapeutic interventions in human congenital toxoplasmosis.

## Introduction

*Toxoplasma gondii* is an obligate intracellular protist that opportunistically infects a wide range of homeothermic animals as intermediate hosts, including humans and small ruminants (1). In humans, *T. gondii* represents a major public health concern due to the risk of transplacental transmission following maternal infection during pregnancy, which can result in abortion, severe fetal development abnormalities, hydrocephalus and long-term neurological and ocular sequelae (2–5). Consequently, early diagnosis is essential to guide clinical management, reduce vertical transmission rates, and prevent severe adverse outcomes for both mother and offspring (2, 3, 6, 7).

Alongside its clinical relevance in human medicine, toxoplasmosis represents a major concern owing to its severe impact on livestock production (1). Similar to humans, primary infections in small ruminants during pregnancy leads to reproductive failure, including fetal loss, abortions or stillbirths, particularly in sheep (1). In fact, *T. gondii* is considered to be linked to 10-23% of ovine abortions in Europe and the US (8). In ovine livestock, early diagnosis is equally critical due to the occurrence of early fetal death (EFD) and early abortions during the second week post-infection (7-14 days pi) (9–12). Under routine field conditions, the early abortions in sheep remain largely unrecognized because conventional diagnostic methods are ineffective during this narrow window of acute phase. Direct parasite identification fails because parasite antigens (immunohistochemistry) and DNA (PCR) remain undetectable in placental and fetal tissues (10, 12). Furthermore, serological testing has limited diagnostic value as maternal immunoglobulin G levels are generally detectable since approximately day 20 pi, after the early abortion occur (13) and IgM (around day 12 pi) suffers from cross-reactivity with other abortifacient pathogens such as *Neospora caninum* (Apicomplexa) and *Chlamydia abortus* (Chlamydiota) (14). In addition, recent findings suggest that this poorly known pathogenic mechanism is hypothetically driven by acute host-derived immunopathological responses, which are strongly influenced by the infective dose (12). However, how infection dose-dependent factors interact with early system immune activation to induce EFD remains to be defined, as only one study has quantified acute serum cytokine levels in sheep experimentally infected with oocysts (15).

To overcome the limitations of conventional diagnostic methods, artificial intelligence and machine learning approaches have emerged as powerful tools capable of identifying non-linear patterns and integrating complex, multidimensional laboratory data and biomarkers to enhance diagnostic accuracy (16, 17). In infectious diseases, machine learning algorithms are increasingly used to optimize early pathogen detection across viral, bacterial and parasitic infections (18, 19). Regarding toxoplasmosis, artificial intelligence has primarily been applied to three areas: detecting retinal lesions in ocular toxoplasmosis (20–22), aiding light microscopy-based examinations to identify parasite’s presence (23), and using white blood cell counts to differentiate between acute and chronic infection (24). However, the integration of circulating host-response biomarkers into machine learning models for the early detection of toxoplasmosis remains unexplored.

Accordingly, the present study aims to characterize the early systemic host response to *T. gondii* infection in pregnant sheep, apply machine learning to identify biomarkers for the early diagnosis of infection, EFD and abortion and gain insights into its underlying pathogenic mechanisms. Beyond addressing these critical veterinary challenges, pregnant sheep represent a translationally superior model for human congenital toxoplasmosis. During acute *T. gondii* infection, the ovine immune response aligns much more closely with human immunopathology that murine models, which suffer from species-specific immune biases that hinder clinical extrapolation (25–28). Therefore, this framework aims to facilitate the translation of both the identified biomarkers and machine learning methodology used into early diagnostic strategies for acute human toxoplasmosis.

## Material and methods

### Ethical issues

All serum samples originated from previously conducted experimental infections in sheep (13, 29–31). Those experiments involving animals were authorized by the Animal Welfare Committee of the Community of Madrid, Spain (PROEX 68.0/20; PROEX 93/14; PROEX 128.1/21; PROEX 128.1/21), following procedures described in the Spanish and European Union legal requirements (Law 32/2007, R.D. 53/2013, and Council Directive 2010/63/EU).

### Experimental design and serum panels

Across all panels, blood samples were collected by jugular venipuncture into 5 mL Vacutainer tubes without anticoagulant (Becton Dickenson and Company, Plymouth, UK). Samples were centrifuged at 450× g for 10 minutes at 4°C to obtain serum samples that were stored at −80°C until analysis.

Serum samples were arranged into six panels **(Table 1)**. All animals used in these experimental infections had been initially confirmed as seronegative for *T. gondii*, *N. caninum*, Border disease virus (BDV), Schmallenberg virus (SBV), *Coxiella burnetii*, and *C. abortus*. Panels 1 and 2 comprised samples from sheep orally inoculated at 90 days of pregnancy with *T. gondii* sporulated oocysts (TgSpSh1, ToxoDB RFLP genotype #3); panel 1 involved animals (*n* = 4) infected with 1,000 oocysts (13) and panel 2 animals (*n* = 18) receiving a dose of 10 oocysts (29, 30). Additionally, panels 1 and 2 included naïve animals (*n* = 6), which serve as negative control. Panel 3 included samples from sheep (*n* = 17) infected at 90 days of pregnancy with different doses of *N. caninum* tachyzoites (31), while panel 4 encompassed samples from pregnant sheep (*n* = 3) infected with different doses of *Chlamydia abortus* (14).

**Table 1:** Serum collection time points and experimental conditions in *Toxoplasma gondii*, *Neospora caninum* and *Chlamydia abortus* experiments.

| Sera panel | Infection dose (pathogen) | Inoculated animals | Non-inoculated animals | Serum collection time points (day pi) | Animals with EFD<sup>5</sup> (day pi) | Reference |
|---|---|---|---|---|---|---|
| 1 | 1,000<br>(*T. gondii*)<sup>1</sup> | 4 | 3 | 0, 2, 3, 4, 5, 6, 7, 8, 9, 10, 14, 21, 28 | 3/4<br>(8 dpi) | (13) |
| 2a<sup>2</sup> | 10<br>(*T. gondii*)<sup>1</sup> | 8 | 3 |  | 1/8<br>(14 dpi) | (30) |
| 2b | 10<br>(*T. gondii*)<sup>1</sup> | 10 | 0 |  | 4/10<br>(10 dpi) | (29) |
| 3a | 10<sup>5</sup><br>(*N. caninum*)<sup>3</sup> | 3 | 0 | 0, 3, 5, 7, 10, 14, 21, 28, 34, 41 | 0/3 | (31) |
| 3b | 10<sup>4</sup><br>(*N. caninum*)<sup>3</sup> | 4 | 0 |  | 0/4 |  |
| 3c | 10<sup>4</sup><br>(*N. caninum*)<sup>4</sup> | 3 | 0 |  | 0/3 |  |
| 3d | 10<sup>3</sup><br>(*N. caninum*)<sup>3</sup> | 4 | 0 |  | 0/4 |  |
| 3e | 10<sup>2</sup><br>(*N. caninum*)<sup>3</sup> | 3 | 0 |  | 0/3 |  |
| 4 | *C. abortus* | 3 | 0 | 15, 30 | 0/3 | (14) |

<sup>1</sup> Oral infection with sporulated oocysts of Type II TgShSp1 *Toxoplasma gondii* isolate (genotype ToxoDB RFLP #3)
<sup>2</sup> In this experimental infection serum samples from 9 dpi were not obtained
<sup>3</sup> Intravenous infections with *Neospora caninum* Nc-Spain 7 isolate tachyzoites
<sup>4</sup> Subcutaneous infections with *Neospora caninum* Nc-Spain 7 isolate tachyzoites
<sup>5</sup>All animals that suffer early fetal death (EFD), expulsed the fetuses (early abortion) 1 or 2 days after.
pi = post-infection

Serum from animals infected under natural conditions has been previously described in a prior study (14) and was expanded in the present work **(Table S1)**. Panel 5 consisted of serum samples from sheep (*n* = 11) that seroconvert to IgG between sampling (11-45 days apart), representing suspected cases of early *T. gondii* infection. Panel 6 included sheep (*n* = 22) from the same flocks that remained seronegative and serve as controls.

The analytical workflow followed in this study is illustrated in **Fig. 1**. Briefly, in step 1, individual host parameters were first analyzed to identify significant differences between high-dose infected sheep (panel 1) and uninfected controls. In step 2, the parameters identified in step 1 were further evaluated in low dose-infected sheep (panel 2) to check dose influence. In step 3, data of all parameters measured in step 2 were incorporated into machine learning analyses to identify the most predictive parameters of early *T. gondii* infection. In step 4, the most predictive parameters determined in step 3 were tested in samples from experimental infections with other abortifacient agents (panel 3 and 4) to replicate the machine learning analyses. Finally, in step 5, the most predictive parameters determined in step 4 were measured in animals exposed under natural conditions (panels 5 and 6) and processed through the machine learning model previously trained in step 4. The parameters analyzed in each panel are detailed in **Supplementary Table S2**.

**Figure 1:**
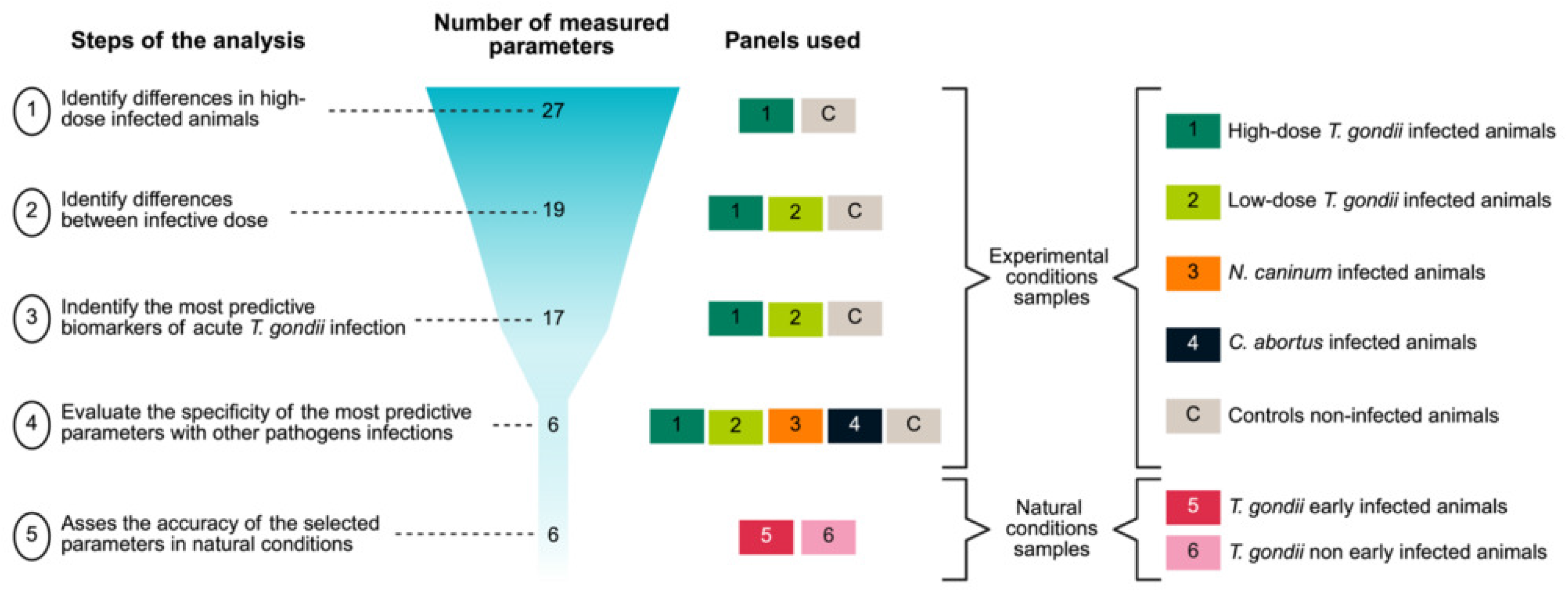
Schematic representation of the study workflow. The sequential analytical steps are shown, indicating the number of host-response parameters evaluated at each stage and the corresponding serum panels used for their assessment. Distribution of serum samples across experimental panels, grouped according to infection type, including experimental *T. gondii*–infected animals, animals experimentally infected with other pathogens associated with reproductive failure, and animals exposed to natural field conditions.

### Analysis of immunological, biochemical, and hemostatic parameters in serum samples

#### Cytokine and enzyme markers of inflammation

Levels of C-X-C motif chemokine ligand 9 (CXCL9), C-X-C motif chemokine ligand 10 (CXCL10), IFN-γ, interleukin 6 (IL-6), interleukin 17 (IL-17) and TNF-α were measured using commercial kits **(Table S3),** based on the standard double-antibody sandwich ELISA technique following the manufacturers’ recommendations. Colorimetric quantification was performed using multi-mode microplate reader (Cytation™ 5, BioTek Instruments, USA).

Adenosine deaminase activity (ADA) (32) and lactate dehydrogenase levels (LDH) (33) were evaluated using an automated biochemistry analyzer (Olympus AU400; Olympus Diagnostica GmbH, Hamburg, Germany).

Total protein and albumin were measured using commercial kits (Total protein OSR 6132, Albumin OSR 6102, Olympus Life and Material Science Europe GmbH, Hamburg, Germany) following the manufactureŕs recommendations with an automated biochemistry analyzer (Olympus AU600; Olympus Diagnostica GmbH, Hamburg, Germany). Globulin concentrations were mathematically calculated as the difference between total protein and albumin values.

### Acute phase proteins quantification

Serum Amyloid A (SAA) concentrations were initially measured for steps 1, 2, and 3 using a latex agglutination turbidimetric immunoassay (LZ-SAA; Eiken® Chemical Co., Ltd.) (34, 35). For the machine learning steps (4 and 5) SAA quantification was shifted to a commercial ELISA kit **(Table S3)**. To ensure consistency across all analyses, all previous serum samples from *T. gondii*-challenged animals were re-measured using this kit, although some samples were no longer available due to insufficient remaining volume (9%). Haptoglobin (Hp) concentrations were determined for all samples using a hemoglobin-binding method with a commercial kit (Tridelta Phase Range Haptoglobin kit; Tridelta Development Limited, Ireland) (36). The initial SAA turbidimetric immunoassay and all haptoglobin determinations were carried out using an automated biochemistry analyzer (Olympus AU600; Olympus Diagnostica GmbH, Hamburg, Germany).

### Estimation of oxidants and antioxidants biomarkers

Oxidative markers included reactive oxygen metabolites (dROMs) (37), total peroxides (POX-Act) (38) and advanced oxidation protein products (AOPP) (39). Antioxidant capacity was evaluated through trolox equivalent antioxidant capacity (TEAC) (40), ferric reducing antioxidant power (FRAP) (41), uric acid (42), cupric reducing antioxidant capacity (CUPRAC) (43), total thiol levels (44) and paraoxonase-1 (PON-1) (45). All determinations were performed using an automated biochemistry analyzer (Olympus AU400; Olympus Diagnostica GmbH, Hamburg, Germany).

### Hemostatic parameters analysis

Blood samples in sodium citrate tubes were sent to Laboratorio de Análisis Veterinarios Arturo Soria S.L. (Madrid, Spain) to measure: prothrombin time (PT) activated partial thromboplastin time (APTT), and fibrinogen using a single-channel optical coagulometer designed for *in vitro* coagulation assays (Coatron M1, Teco, Medical Instruments, Germany).

### Angiopoietin 2 and fibrinogen-like protein 2 quantification

Serum levels of Fibrinogen-like protein 2 (FGL2) and angiopoietin 2 (ANGPT2) were measured using commercial double-antibody sandwich ELISA kits **(Table S3)** following the manufacturers’ recommendations. Colorimetric quantification was performed using multi-mode microplate reader (Cytation™ 5, BioTek Instruments, USA).

### Statistical analyses

As haptoglobin, oxidative stress biomarkers, total proteins and albumin were measured in three different time points, potential batch effect associated with the date of measurement were assessed using principal component analysis (PCA), ANOVA, and linear modeling. A significant influence of the measurement date was observed **(Fig. S1)**, and therefore, the ComBat algorithm implemented in the sva package (version 3.38.0; Bioconductor) within the R environment (version 4.4.3) was applied to adjust for batch effects while preserving biological variability (46).

For all evaluated parameters, statistical time-course analysis was conducted using a mixed-effects model followed by Tukey’s post hoc test. Additionally, when comparing parameters across pathogens (*T. gondii*, *N. caninum*, and *C. abortus*), Kruskal-Wallis test followed by Dunn’s post hoc test was used. All analyses were performed using Python (version 3.12.6) with the scipy.stats and scikit_posthocs packages. Importantly, the SAA concentrations used in the statistical analyses were measured by latex agglutination turbidimetric immunoassay to include all timepoints.

### Machine learning algorithms for predictive data modeling

To comprehensively analyze all parameters together and to explore their potential as predictors of early *T. gondii* infection, we applied a machine learning approach (17). Importantly, the SAA concentrations used in the machine learning were measured by commercial ELISA, as the previously describe assay was unavailable for the other panels included in these analyses (panel 3, 4, 5 and 6).

### Data organization and dataset construction

For the initial predictive modeling, a total of 16 parameters measured in *T. gondii* serum panels 1 and 2 were used: CXCL9, CXCL10, IFN-y, Hp, SAA, ADA, LDH, CUPRAC, FRAP, AOPP, thiol, uric acid, PON-1, TEAC, total proteins and albumin. dROMs and POX-Act were excluded due to excessive missing data, and globulins were omitted because they were derived from total protein and albumin levels. All data were normalized using a min–max scaling.

The normalized data were structured into three different datasets **(Table S4)**. All datasets extended up to 21 dpi, corresponding to the day prior to *T. gondii* IgG seroconversion. Dataset 1 (D1), measurements collected from day 2 pi to capture the whole early phase of infection; dataset 2 (D2), from day 6 pi, when over 90% of infected sheep developed fever (rectal temperature > 40°C); and dataset 3 (D3), from the onset of EFD (days 8-14 pi, depending on the experiment). The control group included non-infected sheep from all days pi and infected sheep at day 0 pi (before inoculation).

To assess the specificity of the models, serum panels from other pathogens experimental infections were include (panel 3 and 4). The infected group included sheep infected with *T. gondii* (panel 1 and 2), while the non-*T. gondii* infected group comprised non-infected sheep (control) and those infected with other pathogens (panels 5 and 6) prior to seroconversion. Two datasets were generated **(Table S4)**: dataset 4 (D4), included all observations from the early phase of infection; and dataset 5 (D5) included observations starting from the onset of fever in each experiment.

### Machine learning algorithms

Three supervised classification algorithms were used to develop the infection predictive models: Decision Tree, Random Forest, and Gradient Boosting (47). All models were implemented in Python (version 3.12.6) using scikit-learn (version 1.5.2) (48) for model training and validation. The class imbalance between groups was addressed by applying class_weight=’balanced’ during model training. For all algorithms, model performance was evaluated using 5-Fold stratified cross-validation and the following classification metrics: accuracy (overall proportion of correct predictions), recall (proportion of correctly identified positives among all actual positives), precision (proportion of correctly predicted positives among all predicted positives) and F1-score (harmonic mean of precision and recall). Hyperparameter tuning was performed using GridSearchCV with customized grids for each model such as maximum tree depth or number of estimators.

To optimize diagnostic efficiency and to identify a minimal set of predictive biomarkers, a feature selection step was implemented using scikit-learn’s “SelectFromModel”. This process was guided by the feature importance scores obtained from the algorithm that exhibited the highest overall performance during the initial evaluation phase. The panel of selected biomarkers identified through this process was subsequently advanced to the multi-pathogen evaluation stage (Datasets 4 and 5). Finally, to validate the clinical utility of the selected panel under natural conditions, raw serum concentrations of the selected biomarkers were first measured in such samples and then normalized using the same min–max scaling previously described. These normalized values were subsequently fed into the pre-trained models to generate the final infection classification results.

### Data availability

The data supporting the findings of this study will be deposited in Dryad Digital Repository, and the corresponding DOI will be provided before publication.

## Results

### High and low-doses of *Toxoplasma gondii* oocyst inoculation trigger a similar proinflammatory response, but a more pronounced redox imbalance was observed in the high-dose inoculated group

Parameters that showed no statistical differences in step 1 were not measured in low-dose infection in step 2, except for total proteins, albumin, SAA, haptoglobin, dROMs, AOPP, FRAP, and PON-1, which showed a trend toward change **(Fig. S2)**.

In step 2, sheep from both *T. gondii*-infected panels (1 and 2) showed significantly elevated levels of several Th1-related parameters IFN-γ, CXCL9, CXCL10 and ADA than control group (*p* < 0.0001) **(Fig. 2).** SAA concentrations were also significantly higher in both panels compared to the control group (*p* < 0.0001) and differenced significantly between panels (*p* < 0.01). In contrast, haptoglobin levels were significantly increased only in panel 2 (low-dose group) (*p* < 0.0001) **(Fig. 2)**. On the other hand, oxidative stress was more pronounced in panel 1 (high-dose group), with significantly higher dROMs levels than both the control group and panel 2 (low-dose group) (*p* < 0.0001) and lower antioxidant capacity. TEAC and CUPRAC levels were significantly lower in panel 1 (high-dose group) compared to control (*p* < 0.0001 and *p* < 0.01, respectively) and panel 2 (low-dose group) (*p* < 0.0001 for both), while FRAP concentrations were significantly reduced in panel 1 (high-dose group) compared to control (*p* < 0.05). Conversely, uric acid levels significantly increased in both panels compared to control (*p* < 0.01). Additionally, PON-1 and total protein levels were significantly increased in panel 1 (high-dose group) compared to controls (*p* < 0.05). In contrast, globulin concentrations exhibited a significant increase in panel 2 (low-dose group) compared to the control group (*p* < 0.05) **(Fig. 2).**

**Figure 2:**
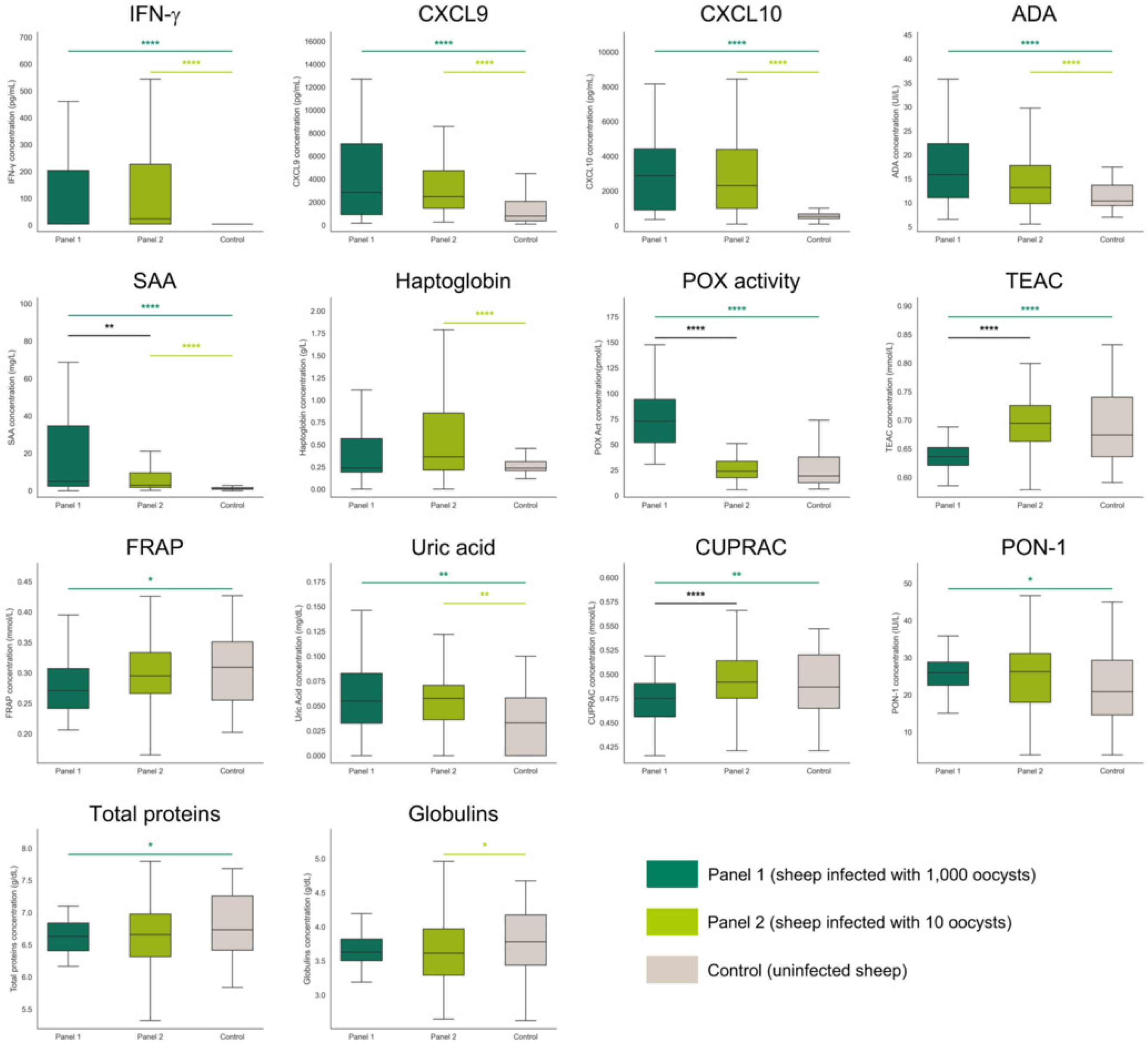
Differential cytokine and biomarker profiles in response to *Toxoplasma gondii* infection at different doses of oocysts. Experimental groups are color-coded as follows: Panel 1 (dark green), Panel 2 (light green), and control (grey). Bar plots show molecules with statistically significant differences among experimental groups (Panel 1, Panel 2, and control). The SAA concentrations shown were determined using the latex agglutination turbidimetric immunoassay. Statistical analysis was performed using a mixed-effect model followed by Tukey’s multiple comparisons test between groups. Boxes represent the interquartile range (IQR) with the median indicated by the central line; whiskers extend to 1.5×IQR. Statistical significance is indicated as \**p* < 0.05, \*\**p* < 0.01, \*\*\**p* < 0.001 and \*\*\*\**p* < 0.0001 using different colors: dark green for differences between panel 1 and control, light green for differences between panel 2 and control and black for differences between panels.

The graphical representation of the temporal dynamics of each of these parameters can be found in Fig. S2.

### Marked oxidative stress and serum amyloid A elevation are hallmarks of early fetal death during ovine congenital toxoplasmosis

Serum panels from panels 1 and 2 were re-analysis according to reproductive outcomes **(Table 1)**. Only parameters exhibiting significant differences between sheep that suffer EFD and subsequent abortion and those that maintained pregnancy are presented in **Fig. 3**. Importantly, all animals that experienced EFD episodes, expulsed the fetuses (early abortion) within 1 or 2 days of fetal death.

**Figure 3:**
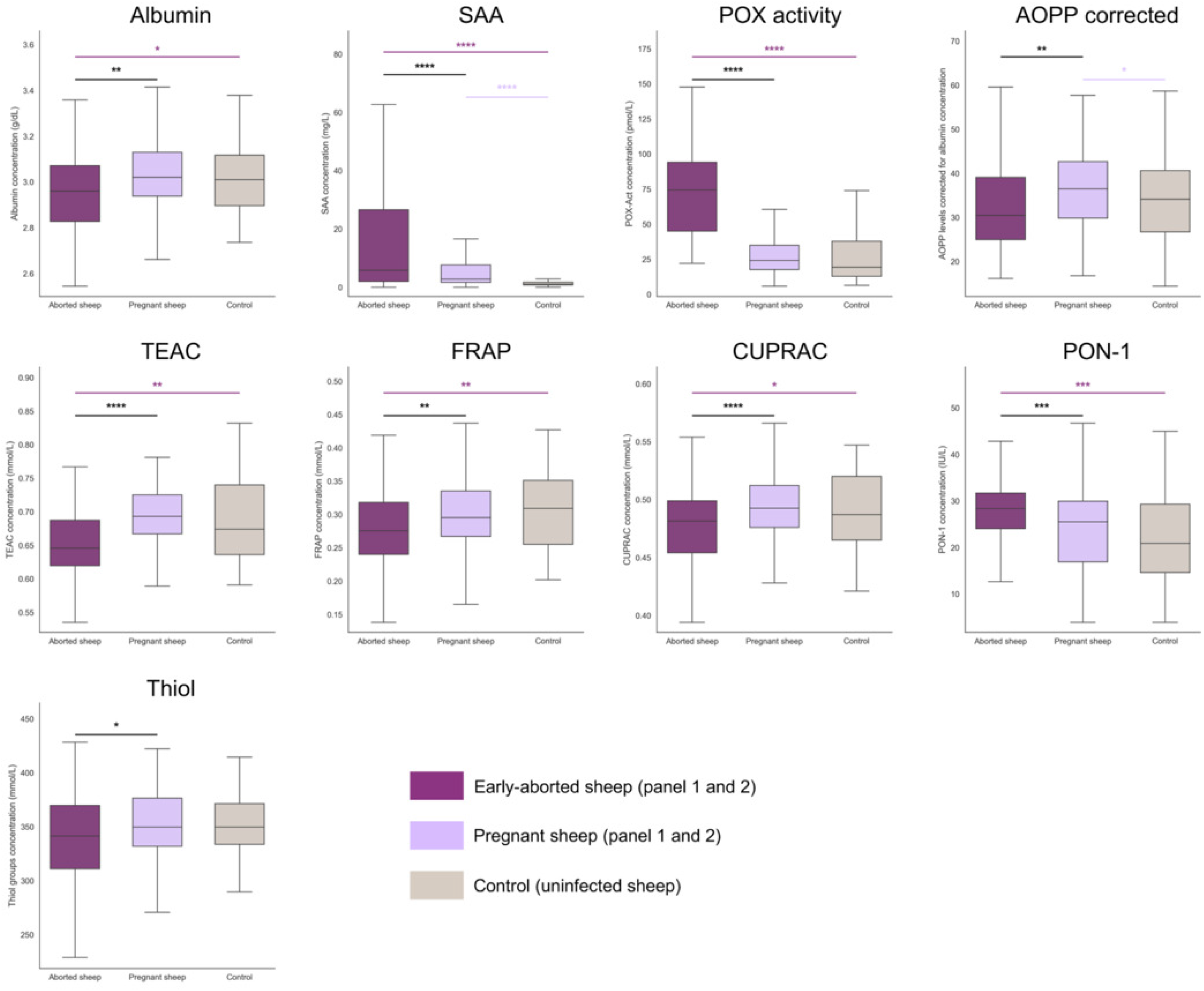
Differential cytokine and biomarker profiles associated with pregnancy status and early fetal death outcomes following *Toxoplasma gondii* infection. Experimental groups are color-coded as follows: animals from panel 1 and 2 that suffer for early fetal death and abortion (dark purple), animals from panels 1 and 2 that maintained pregnancy (light purple), and control non-infected animals (grey). Boxplots show molecules with statistically significant differences between pregnant and aborted animals. The SAA concentrations shown were determined using the latex agglutination turbidimetric immunoassay. Statistical analysis was performed using a mixed-effect model followed by Tukey’s multiple comparisons test between groups. Boxes show the interquartile range (IQR) and the median; whiskers extend to 1.5×IQR. Statistical significance is indicated as \**p* < 0.05, \*\**p* < 0.01, \*\*\**p* < 0.001 and \*\*\*\**p* < 0.0001 using different colors: dark purple for differences between aborted animals and control, light purple for differences between pregnant animals and control and black for differences between aborted and pregnant infected animals.

In this context, albumin concentrations showed different levels in both groups, being significantly lower in the aborted group compared to control (*p* < 0.05) and compared to non-aborted group (*p* < 0.01). The acute phase protein SAA was significantly elevated in both aborted and non-aborted groups relative to control (*p* < 0.0001); however, concentrations were significantly higher in the aborted group than in the non-aborted group (*p* < 0.0001). This dynamic was noticeable from day 6 pi, when the aborted group exhibited a higher increase over controls than the non-aborted group **(Fig. S3)**. Oxidative stress markers showed the most striking differences related to the reproductive outcome **(Fig. 3)**. The aborted group exhibited severe oxidative stress, with POX activity being significantly higher than both the control group (*p* < 0.0001) and the non-aborted group (*p* < 0.0001). However, AOPP corrected with albumin levels were significantly elevated in the non-aborted group compared to the aborted group (*p* < 0.01) and the controls (*p* < 0.05). Antioxidant capacity was notably depleted in sheep that suffered EFD, with lower levels of TEAC, FRAP (*p* < 0.01) and CUPRAC (*p* < 0.05) compared to control animals and with non-aborted animals (*p* < 0.0001 for CUPRAC and TEAC and *p* < 0.01 for FRAP). Conversely, PON-1 activity showed an increase in aborted animals, being significantly higher compared to control and to non-aborted group (*p* < 0.001). Kinetic analysis revealed that the surge in oxidant markers occurred near the EFD episode, with significant increases in POX activity in the aborted group starting at day 6 pi, with significant changes started at day 10 pi (*p* < 0.05). In contrast, the decline in antioxidant defenses occurred later; CUPRAC and TEAC levels began to decrease significantly at days 8-10 pi, followed by a significant drop in total thiols from day 14 pi onwards. However, FRAP levels start to decrease in the aborted group as early as day 6 pi. **(Fig. S3)**

The plots representing the temporal dynamics of each of these parameters can be found in **Fig. S3**.

### Random Forest algorithm identifies a reduced panel of four predictive biomarkers, achieving high performance in differentiating *T. gondii*-infected animals from non-infected controls

For the predictive modeling in step 3, the serum biomarker data from *T. gondii* infected sheep (panels 1 and 2) were compiled, normalized and organized into three time-dependent datasets (D1–D3).

As shown in **Table 2**, all three algorithms used reached accuracies of 71-76% in D1 (from day 2 pi, early phase), increasing in D2 (from day 6 pi, fever onset) and D3 (from days 8-14 pi, fetal death onset) above 94%, with balanced precision, recall and F1-scores.

**Table 2.** Performance comparison of Decision Tree, Random Forest and Gradient Boosting models across five datasets, including the impact of Select From Model (SFM) thresholding on final classification metrics.

| Dataset | Machine learning algorithm | Threshold | Parameters selected<sup>1</sup> | Accuracy<sup>3</sup> | Precision<sup>4</sup> | Recall<sup>5</sup> | F1-Score<sup>6</sup> |
|---|---|---|---|---|---|---|---|
| 1 | DT |  |  | 71% | 76% | 71% | 73% |
|  | RF |  |  | 76% | 80% | 76% | 77% |
|  | RF-SFM<sup>1</sup> | 0.05 | CXCL10, IFN-γ, SAA<sup>2</sup> | 65% | 66% | 65% | 66% |
|  | GB |  |  | 72% | 75% | 72% | 73% |
| 2 | DT |  |  | 95% | 95% | 95% | 95% |
|  | RF |  |  | 94% | 94% | 94% | 94% |
|  | RF-SFM<sup>1</sup> | 0.05 | CXCL9, CXCL10, SAA<sup>2</sup> | 87% | 87% | 87% | 87% |
|  | GB |  |  | 97% | 97% | 97% | 97% |
| 3 | DT |  |  | 91% | 91% | 91% | 91% |
|  | RF |  |  | 89% | 89% | 89% | 89% |
|  | RF-SFM<sup>1</sup> | 0.1 | CXCL10, SAA<sup>2</sup> | 89% | 90% | 89% | 89% |
|  | GB |  |  | 95% | 95% | 95% | 95% |
| 4 | DT |  |  | 74% | 74% | 74% | 74% |
|  | RF |  |  | 78% | 79% | 79% | 78% |
|  | GB |  |  | 78% | 78% | 78% | 78% |
| 5 | DT |  |  | 78% | 79% | 78% | 78% |
|  | RF |  |  | 80% | 80% | 80% | 80% |
|  | GB |  |  | 82% | 82% | 82% | 82% |

<sup>1</sup> Features were selected using “SelectFromModel” according to the importance scores of the best-performing initial model
<sup>2</sup> SAA concentrations used were measured using the commercial ELISA
<sup>3</sup> Overall proportion of correct predictions
<sup>4</sup> Proportion of correctly predicted positives among all predicted positives
<sup>5</sup> Proportion of correctly identified positives among all actual positives
<sup>6</sup> Harmonic mean of precision and recall
DT = Decision Tree
RF = Random Forest
RF-SFM = Random Forest - SelectFromModel
GB = Gradient Boosting

Although Gradient Boosting performed slightly better in D2 (from day 6 pi, fever onset) and D3 (from days 8-14 pi, fetal death onset), Random Forest reached the best performance in D1, which represents the early phase of infection, the most vital stage for timely diagnosis, before the onset of fever (D2) or abortion (D3). Therefore, the most predictive parameters selection was performed using Random Forest and scikit-learn’s “SelectFromModel”. Within this framework, each parameter is assigned to a specific weight (importance score) by the algorithm. By increasing the selection threshold from its default baseline of zero, we can restrict the model to utilize only the most dominant parameters, maintaining a balance between high classification performance (accuracy, precision, recall, F1-score) and identifying a minimal, realistic set of biomarkers. Based on this, a threshold of 0.05 was selected for D1 and D2, while a stricter threshold of 0.1 was applied to D3 **(Table 2)**. Consequently, the selected parameters were: CXCL10, IFN-y, and SAA for D1; CXCL9, CXCL10, and SAA for D2 and CXCL10 and SAA for D3. Taken together, CXCL10, CXCL9, IFN-y and SAA were selected and incorporated into datasets D4 and D5 to evaluate the specificity of these parameters against *N. caninum* (panel 3) and *C. abortus* (panel 4) infections in the step 4.

### Decision Tree identifies key discriminatory biomarkers in multi-pathogen datasets with moderate overall performance

Before retraining the machine learning models (step 4), we analyzed the baseline statistical differences in cytokine biomarker levels across the distinct panels **(Fig. S4).** This analysis revealed that CXCL9 levels were significantly higher in the *T. gondii*-infected group (panels 1 and 2) compared to both *N. caninum* (panel 3) and *C. abortus* (panel 4) (*p* < 0.001) infected sheep. Conversely, SAA concentrations were markedly elevated in the *N. caninum* group (panel 3) compared to *T. gondii* (panels 1 and 2) (*p* < 0.001) and *C. abortus* (panel 4) (*p* < 0.01).

Interestingly, in D4 (early infection phase), predictive performance improved markedly, with metrics across all algorithms ranging between 74% and 78% for accuracy, precision, recall, and F1-scores. Conversely, in D5 (from fever onset), the multi-pathogen complexity led to a slight reduction in overall performance compared to the previous binary stage, although accuracies remained stable around 78– 82% with concordant values across all other metrics **(Table 2)**.

Finally, all algorithms trained on dataset D4 (early infection phase) were selected to evaluate their diagnostic performance under field conditions (step 5) using natural infection-derived samples (panels 5 and 6). Dataset D4 (early infection phase) was prioritized over D5 (from fever onset) for this validation stage because rectal temperature is expected to be unknown in field scenarios.

### Integration of host biomarkers and predictive algorithms successfully bridges the diagnostic gap in natural abortion outbreaks of ovine toxoplasmosis

In the final step of the analytical workflow (step 5), the diagnostic performance of the selected biomarkers was validated under field conditions using serum panels 5 (early *T. gondii*-infected animals) and 6 (control animals). Serum concentrations of IFN-y, CXCL9, and CXCL10 from the first sample point were determined using commercial ELISA kits **(Table S3)**, normalized using the original min-max scaling protocol, and then processed through the Decision Tree, Random Forest, and Gradient Boosting models pre-trained in step 4.

Among the models tested, the Decision Tree **(Fig. 4)** yielded the lowest error rate, achieving an overall accuracy of 84.84% **(Table 3)**. Notably, this model demonstrated a sensitivity of 100%, successfully identifying every sheep suffering early *T. gondii* infection from panel 5 (early *T. gondii*-infected animals). The specificity was 77.27%, as the model classified 5/22 control animals from panel 6 (control animals) as infected.

**Figure 4:**
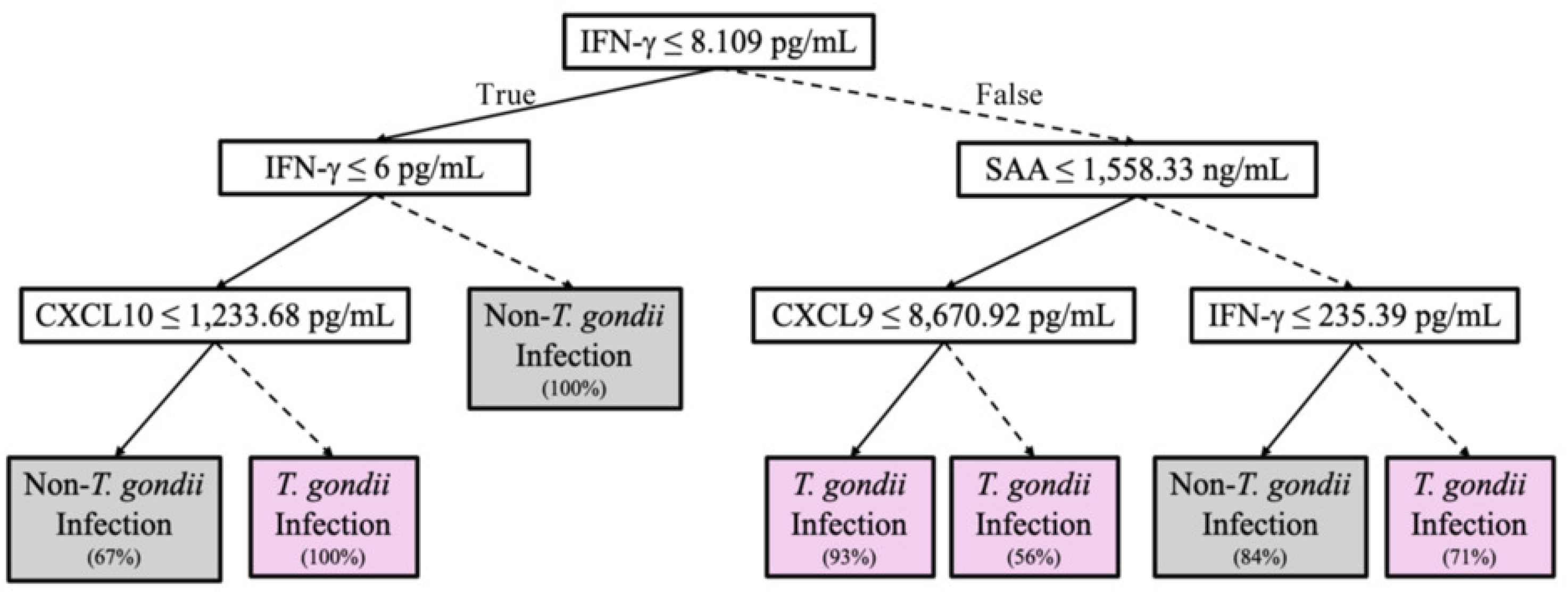
Structure of the final Decision Tree algorithm for the prediction of a *Toxoplasma gondii* infection compared to other pathogen infections and non-infected animals during the early stage of infection (Dataset 4). The model is interpreted from top to bottom. Each internal node represents a decision rule based on a biomarker threshold. If the condition is true, the left branch (dashed line) is followed; if false, the right branch (solid line) is followed. Terminal nodes indicate the predicted class: “*T. gondii”* (samples classified as animals infected with *T. gondii*) or “Non-*T. gondii”* (animals either uninfected or infected with other agents: *Neospora caninum* or *Chlamydia abortus*). The percentage shown in each terminal node represents the proportion of correctly classified samples reaching that node.

**Table 3:** Diagnostic performance and field validation of previously trained *Machine Learning* models (Decision Tree, Random Forest and Gradient Boosting) for early *T. gondii* detection in sheep.

| Sera panel | Animal ID | Pregnancy status | DT classification | RF classification | GB classification |
|---|---|---|---|---|---|
| Panel 5<br>(early *T. gondii*-infected animals) | 1 | Aborted | *T. gondii* | *T. gondii* | *T. gondii* |
|  | 2 | Unknown | *T. gondii* | *T. gondii* | *T. gondii* |
|  | 3 | Unknown | *T. gondii* | *T. gondii* | *T. gondii* |
|  | 4 | Unknown | *T. gondii* | *T. gondii* | *T. gondii* |
|  | 5 | Unknown | *T. gondii* | Non-infected | Non-infected |
|  | 6 | Unknown | *T. gondii* | *T. gondii* | *T. gondii* |
|  | 7 | Aborted | *T. gondii* | *T. gondii* | *T. gondii* |
|  | 8 | Aborted | *T. gondii* | *T. gondii* | *T. gondii* |
|  | 9 | Pregnant | *T. gondii* | *T. gondii* | *T. gondii* |
|  | 10 | Pregnant | *T. gondii* | *T. gondii* | *T. gondii* |
|  | 11 | Pregnant | *T. gondii* | *T. gondii* | *T. gondii* |
| Panel 6<br>(control animals) | 1 | Aborted | *T. gondii* | *T. gondii* | Non-infected |
|  | 2 | Aborted | Non-infected | *T. gondii* | Non-infected |
|  | 3 | Aborted | Non-infected | Non-infected | Non-infected |
|  | 4 | Pregnant | Non-infected | Non-infected | Non-infected |
|  | 5 | Pregnant | Non-infected | Non-infected | Non-infected |
|  | 6 | Unknown | Non-infected | Non-infected | Non-infected |
|  | 7 | Unknown | Non-infected | Non-infected | Non-infected |
|  | 8 | Unknown | Non-infected | Non-infected | Non-infected |
|  | 9 | Unknown | Non-infected | Non-infected | Non-infected |
|  | 10 | Pregnant | Non-infected | Non-infected | Non-infected |
|  | 11 | Pregnant | *T. gondii* | *T. gondii* | *T. gondii* |
|  | 12 | Pregnant | Non-infected | Non-infected | Non-infected |
|  | 13 | Pregnant | Non-infected | Non-infected | Non-infected |
|  | 14 | Pregnant | Non-infected | Non-infected | Non-infected |
|  | 15 | Pregnant | Non-infected | Non-infected | Non-infected |
|  | 16 | Pregnant | *T. gondii* | *T. gondii* | *T. gondii* |
|  | 17 | Pregnant | Non-infected | Non-infected | Non-infected |
|  | 18 | Pregnant | *T. gondii* | *T. gondii* | *T. gondii* |
|  | 19 | Pregnant | Non-infected | Non-infected | Non-infected |
|  | 20 | Pregnant | *T. gondii* | *T. gondii* | *T. gondii* |
|  | 21 | Pregnant | Non-infected | Non-infected | Non-infected |
|  | 22 | Pregnant | Non-infected | Non-infected | Non-infected |
| Accuracy<sup>1</sup> |  |  | 84.84% | 78.78% | 84.84% |
| Sensitivity<sup>2</sup> |  |  | 100% | 90.9% | 90.9% |
| Specificity<sup>3</sup> |  |  | 77.27% | 72.72% | 81.81% |

Non-infected = non-*T. gondii* infection
<sup>1</sup>Overall proportion of correct predictions
<sup>2</sup>Proportion of actual positive cases that are correctly identified as positive by the algorithm
<sup>3</sup>Proportion of actual negative cases that are correctly identified as negative by the model
DT = Decision Tree
RF = Random Forest
GB = Gradient Boosting

## Discussion

Despite the increasing recognition of EFD as a distinct presentation of ovine congenital toxoplasmosis (12), its diagnosis is challenging and its underlying pathogenic mechanism remains poorly understood (10, 11, 13). Although their characteristics suggest host-mediated processes, the systemic immune response during this phase has been insufficiently studied, limiting the identification of biomarkers and consequently the understanding of how the infective dose, the primary variable influencing the occurrence of EFD (12), modulates this early systemic response. In this context, despite the tenfold difference in inoculum, the kinetics and magnitude of IFN-y, CXCL9 and CXCL10 responses were similar, indicating that even low-dose exposure is sufficient to trigger a robust Th1-type response. The early rise in peripheral IFN-γ aligns previous observations with 50 sporulated oocysts (day 5 pi) (15). This cytokine induces the production of the chemokines CXCL9 and CXCL10 (49, 50), crucial for the T-cells recruitment, amplifying the adaptive immune response and limiting parasitic dissemination (50).

Parallel to this cellular signaling, the systemic inflammatory cascade was also triggered, as evidenced by an early increase of serum amyloid A (SAA) and haptoglobin in *T. gondii*-infected sheep. These findings are consistent with previous studies of *T. gondii* infection in mice (51, 52), and pigs (53). In our study, the significant SAA elevations in aborted sheep compared to pregnant ones on days 10 and 14 pi aligned closely with EFD. Beyond being an inflammatory marker, SAA promotes platelet aggregation, enhances amyloid incorporation into fibrin, and induces tissue factor expression in endothelial cells, directly initiating the coagulation cascade (54, 55). Thus, early and sustained SAA elevation in aborted sheep likely reflects a hypercoagulability and inflammatory state that may drives EFD.

Beyond inflammation, host control of *T. gondii* relies on reactive oxygen (ROS) and nitrogen species (RNS) (28, 56, 57). However, chemokine and IFN-y driven ROS generation can overwhelm endogenous antioxidant defenses (58). In our study, we observed this oxidative imbalance, defined as oxidative stress, in sheep infected with a high dose of *T. gondii* oocysts. This oxidative stress occurs across host species including mice (59), rats (60, 61), naturally infected sheep (62) and humans (63). Importantly, this severe redox imbalance was significantly more pronounced in those that suffered EFD and abortions, characterized by placental thrombosis. In this context, oxidative stress has been increasingly recognized as a critical driver of coagulation and thrombosis by promoting thrombin generation, thrombus growth and stability (64, 65) and a hypercoagulable state (66). Furthermore, the redox imbalance damages placental cells (67, 68), promoting endothelial dysfunction by reducing nitric oxide bioavailability, disrupting angiogenic signaling, and ultimately restricting maternal-fetal nutrient and oxygen transfer (69–71). Such pathological oxidative stress is a known driver of gestational disorders, including preeclampsia, fetal growth restriction, and recurrent miscarriage (72–74). In our study, sheep without EFD exhibited significantly lower redox alterations regardless of dose, demonstrating that oxidative stress intensity-rather than the infective dose itself drives thrombosis and placental damage during ovine toxoplasmosis.

The multifaceted nature of these responses, suggest that the host-pathogen interaction is too intricate to be fully interpreted through single-parameter analyses. Many studies investigated individual biomarkers for *T. gondii* infections in animals (75–78) but little effort has been paid to integrate these assorted host-response-related parameters in a unified model. In this regard, machine learning provides a powerful framework precisely for such integrative, multivariate analysis (16). In our study, the Decision Tree model based in four biomarkers (CXCL9, CXCL10, IFN-y and SAA) exhibited a remarkable diagnostic sensitivity of 100%, successfully identifying all positive cases during the early-infection window. From a clinical perspective, the validation of this refined 4-biomarker panel using field serum samples opens the door for the development of targeted diagnostic assays and provides an approach to address the difficulty of early diagnosis in human toxoplasmosis.

The diagnostic and clinical relevance of these four mediators is well supported by prior studies carried out in humans. The chemokines CXCL9 and CXCL10 have been identified as key indicators of systemic inflammation and ocular commitment; CXCL9 serves as serum hallmark for active retinechoroidal lesions in infants with congenital toxoplasmosis (79), whereas CXCL10 and CXCL9 combined allow for the stratification of infection phases and ocular involvement in serum (80), as well as in asymptomatic seropositive donors (81), and in intraocular fluid from patients with ocular toxoplasmosis (82). Furthermore, IFN-y directly correlate with active retinechoroidal damage in congenitally infected infants (79, 83) as well as in patients with primary acquired recurrent ocular toxoplasmosis (82). Finally, serum amyloid A (SAA) has been characterized in infected adults as a systemic biomarker reflecting chronic inflammation and vascular reactivity (84). Together, these preceding human findings reinforce the biological plausibility and clinical utility of integrating these four specific factors into a unified diagnostic panel.

Compared with conventional serology, which is hindered by delayed IgG seroconversion and long-persisting IgM (85, 86), this machine learning framework enables early diagnosis from a single blood sample. To date, artificial intelligence applications in toxoplasmosis relied on image analysis (20–23). Here, integrating a minimal four-biomarkers panel effectively captures non-linear host responses (16, 17) offering a novel approach to the early diagnosis of toxoplasmosis.

In conclusion, from a veterinary medicine point of view, this 4-biomarkers panel provides a promising tool for early disease detection and the identification of previously unrecognized EFD and early abortion episodes in sheep flocks. Beyond its diagnostic utility, our findings reveal that severe oxidative stress may act as the key pathogenic drivers of placental damage and EFD, offering essential immunopathogenic insights into ovine toxoplasmosis. On the other hand, translating this host-signature strategy to human medicine represents a remarkable step forward. Evaluating these biomarkers in pregnant women during early acute infection, aiming a similar validation approach established here, could bridge the current diagnostic gap and facilitate timely clinical interventions to prevent congenital toxoplasmosis and gestational loss in women.

## Acknowledgments

This study was funded by the Spanish Ministry of Science and Innovation (Project PID2022-138673OB-C21, funded by MICIU/AEI/ 10.13039/501100011033 and by “ERDF/EU”) and supported by the Community of Madrid (TEC-2024/BIO-66/SALAINDEC-CM). Ana Huertas-López is the recipient of a Margarita Salas postdoctoral fellowship at the University of Murcia, awarded under the Spanish University System Requalification Programme (Spanish Ministry of Universities) and funded by the European Union-NextGenerationEU (R-1593/2022). Natalia Velasco Jiménez is the recipient of a predoctoral contract (Ministry of Science, Innovation and universities, Spanish State Research Agency; PREP2022-000438). We are particularly grateful to Dr. Andrea Largo de la Torre and Dr. Javier Regidor Cerrillo (SALUVET-INNOVA) for generously providing serum samples, and to Lucía Magal Ranchal for her excellent technical assistance.

The funders had no role in study design, data collection and interpretation, or the decision to submit the work for publication.

NVJ: Data curation, Formal analysis, Investigation, Methodology, Visualization, Writing – original draft, Writing – review & editing; RSS: Formal analysis, Investigation, Methodology, Writing – review & editing; AHL: Formal analysis, Investigation, Methodology, Writing – review & editing; CPR: Methodology, Writing – review & editing; FHL: Software, Methodology, Validation, Writing – review & editing; JB: Investigation, Writing – review & editing; DGE: Investigation, Writing – review & editing; JJC: Writing – review & editing; PHI: Conceptualization, Investigation, Writing – review & editing; RCB: Conceptualization, Investigation, Funding acquisition, Writing – review & editing; LOM: Conceptualization, Investigation, Funding acquisition, Writing – review & editing

## Supplementary material

**Figure S1.** Principal Component Analysis (PCA) of experimental measurement data before and after the batch effect correction.

**Figure S2.** Kinetics of immune and acute-phase biomarkers in panel 1, 2 and controls.

**Figure S3.** Kinetics of immune and acute-phase biomarkers in aborted and non-aborted sheep.

**Figure S4.** Biomarkers levels in panels 1 and 2 (*T. gondii*), 3 (*N. caninum*) and 4 (*C. abortus*)

**Table S1.** Individual serological results, sampling intervals and pregnancy status of sheep during natural *Toxoplasma gondii* outbreaks (panel 5 and 6).

**Table S2.** Host response parameters specifically evaluated in each experimental sera panel

**Table S3.** Information about commercial ELISA kits used for quantification of biomarkers

**Table S4.** Characteristics of all datasets used for the machine learning model

